# Clinically relevant orthotopic pancreatic cancer models for adoptive T cell therapy

**DOI:** 10.1101/2023.09.07.556707

**Authors:** Natalie K. Horvat, Isaac Karpovsky, Maggie Phillips, Megan Wyatt, Margaret A. Hall, Cameron Herting, Jacklyn Hammons, Zaid Mahdi, Richard A. Moffitt, Chrystal M. Paulos, Gregory B. Lesinski

**Affiliations:** Department of Hematology and Oncology, Winship Cancer Institute of Emory University, Atlanta, GA, USA; Department of Surgery, Winship Cancer Institute of Emory University, Atlanta, GA, USA; Department of Pathology and Laboratory Medicine, Winship Cancer Institute of Emory University, Atlanta, GA, USA

**Keywords:** PDAC, ACT, mouse, GEMM, desmoplasia, mesothelin, p53

## Abstract

Pancreatic ductal adenocarcinoma (PDAC) is an aggressive tumor. Prognosis is poor and survival is low in patients diagnosed with this disease; ∼12% at 5 years. Immunotherapy, including adoptive T cell transfer therapy, has not impacted outcomes in PDAC patients, due in part to the hostile tumor microenvironment (TME) which limits T cell trafficking and persistence. We posit that murine models serve as useful tools to study the fate of T cell therapy. Currently, genetically engineered mouse models (GEMM) for PDAC are considered a “gold-standard” as they recapitulate many aspects of human disease. However, these models have limitations, including marked tumor variability across individual mice and cost of colony maintenance. We characterized the immunologic features and trafficking patterns of adoptively transferred T cells in orthotopic PDAC models using two mouse cell lines, KPC-Luc and MT-5, isolated from KPC-GEMM mouse models (Kras^LSL-G12D/+^p53^-/-^ and Kras^LSL-G12D/+^p53^LSL-R172H/+^, respectively). The MT-5 orthotopic model best recapitulates the cellular and stromal features of the TME in the PDAC GEMM. In contrast, far more host immune cells infiltrate KPC-Luc tumors, which have less stroma. Albeit CD4+ T cells were similarly detected in MT-5 tumors compared to KPC-GEMM in mice. Interestingly, we found that CAR T cells redirected to recognize mesothelin on these tumors that signal via CD3ζ and 41BB (Meso-41BBζ-CAR T cells) post antigen recognition infiltrated the tumors of mice bearing stroma-devoid KPC-Luc orthotopic tumors, but not MT-5 tumors. Our data establish for the first time a reproducible and realistic clinical system useful for modeling stroma-rich and stroma-devoid PDAC tumors. These models shall serve in-depth study of how to overcome barriers that limit anti-tumor activity of adoptively transferred T cells.

## Introduction

Pancreatic ductal adenocarcinoma (PDAC) is a deadly malignancy. Only 12% of patients experience an overall 5-year survival rate. PDAC is projected to become the second-leading cause of cancer-related death by 2030^1,2^. With limited therapeutic options, surgery remains the only curative approach. Unfortunately, early-stage disease is usually not detected, resulting in most individuals (80% of patients) being diagnosed with locally advanced or metastatic disease. This diagnosis renders them ineligible for surgical resection^3^. Current standard of care includes cytotoxic chemotherapies, such as FOLFIRINOX or gemcitabine and nab-paclitaxel, which result in only incremental improvements to survival^4^. Similarly, immunotherapy has not impacted disease course, aside from rare subsets of patients harboring microsatellite instable tumors^5^. It is dire to better illuminate the complex biology of PDAC and support efforts in developing new immune-based therapeutics for patients.

PDAC harbors a tumor microenvironment (TME) consisting of abundant non-neoplastic cells that crosstalk with PDAC tumors to produce a dense, desmoplastic stroma. Stroma accounts for ∼90% of tumors by mass, and the degree of desmoplasia in PDAC tumors is linked to limited efficacy of cytotoxic chemotherapy and negative survival outcomes^6^. Prior efforts to target desmoplasia using broad acting agents such as PEG-hyaluronidase have not been met with clinical success^7,8^, prompting the need to investigate the complex biology of the TME. In PDAC, the stroma is fueled by cancer-associated fibroblasts (CAFs), which produce extracellular matrix (ECM) constituents, such as collagen, to increase fibrosis^9^. Adding to this complexity is the heterogeneous origin and function of CAFs, which have context-dependent roles in shaping the PDAC TME and response to therapy^10–13^. Notably, CAFs produce a multitude of soluble factors that influence immune cell composition in PDAC tumors, and compromise the infiltration, survival, and function of cytotoxic lymphocytes^14–16^. Combining these immune features with a low tumor mutation burden, and elevated inhibitory immune checkpoint ligands, PDAC is considered an “immune desert”^17,18^. Taken together, the dominant stromal features of PDAC limit vascularization, access of pharmacologic agents, and infiltration of both endogenous or adoptively transferred immune cells into tumors^19^.

Identifying effective strategies to overcome these redundant stromal barriers could have marked impact, particularly in applying immunotherapy to PDAC. This idea is especially relevant for development of cellular therapies in advanced disease. Efficacy of adoptive T cell transfer (ACT) therapy in PDAC is hindered by numerous factors, including poor persistence and limited access or trafficking of transferred T cells into tumors. For instance, a Phase I clinical trial treated patients with lentiviral-encoded mesothelin-specific chimeric antigen receptor (CAR) T cells and conferred stable disease in 11/15 patients with PDAC, ovarian carcinoma, or malignant pleural mesothelioma. Yet efficacy was limited, likely due to poor infiltration and persistence of immune cells in the TME^20^. ACT therapy studies with CAR T cells targeting other distinct tumor antigens have produced similar results, marked by nominal access of T cells into tumors^21^. Therefore, the ability to accurately model the cellular composition of the TME and its interplay with the immune system is crucial for developing effective therapies for PDAC.

We posit that murine models are useful surrogates for accelerating pre-clinical efforts to improve PDAC therapy. Historically, genetically-engineered mouse models (GEMM) with oncogenic *Kras* and alterations in *tp53,* including those with hot spot, loss of function mutations, or others completely null for *tp53.* Collectively, these GEMM have informed our understanding of the TME and been dependable tools for studying the complex aspects of this disease in an immune competent setting. However, these ‘gold-standard’ GEMM have obvious limitations for rapid therapeutic studies including cost, variable disease onset, and extended time to tumor development. Over time, these GEMM have given rise to useful syngeneic cell lines that can be orthotopically implanted into the pancreas of mice to generate organotropic tumors. These derivative models are not in and of themselves spontaneously arising but do offer the advantage of rapidly generating large cohorts of tumor-bearing mice to minimize variable disease onset for pre-clinical studies. Although derivative cell lines are genetically very similar, they often possess variability in progression. These data imply that subtle differences in the resulting TME may occur between models, and that these differences could be exploited for approximating PDAC stromal heterogeneity. We hypothesized that syngeneic PDAC tumor lines produce distinct stromal features when grown orthotopically *in vivo,* which in turn impact access of immune cells to infiltrate tumors. We found that distinct stromal features of PDAC tumor models can differentially influence T cell trafficking and function in the context of ACT therapy.

## Materials and methods

### Cell Lines and Reagents

KPC-Luc cells (a gift from Dr. Craig Logsdon, MD Anderson Cancer Center) were derived from the Kras^LSL-G12D^, p53^-/-^, Pdx1-cre spontaneous tumor model and express an enhanced firefly luciferase construct ^22^. Cells were cultured with DMEM (Gibco #11965092) with 10% FBS (GeminiBio #900-208), 10 nM L-glutamine (Corning #25-005CI) and Antibiotic:Antimycotic Solution (GeminiBio #50-753-3020). MT-5 cells, derived from the Kras^LSL-G12D^, Trp53^LSL-R270H^, Pdx1-cre spontaneous tumor model were a gift from Dr. David Tuveson (Cold Spring Harbor Laboratory, Cold Spring Harbor, NY) and were cultured with RPMI-1640 (Gibco# 11875093) with 10% FBS, 10 mM L-glutamine, and Antibiotic:Antimycotic Solution. MT-5 cell line were also transduced to stably express luciferase using Lentifect^TM^ (GeneCopoeia #LPP-HLUC-Lv105-025-C) lentiviral vectors of firefly luciferase, according to the manufacturer’s protocol. Single-cell clones of transduced cells were selected and expanded for *in vivo* validation. Briefly, cell populations were diluted to 0.5 cells per 200 µL and plated into a 96-well plate. Each single-cell colony was transferred to T25 flasks and allowed to passage several times to allow for stable colony formation. Cells were then subcutaneously injected into the flank of C57BL/6 mice. Bioluminescence intensity was measured using the IVIS system (described below). Colonies with detectable luciferase expression that displayed longitudinal growth subcutaneously were then implanted orthotopically in the pancreas of mice for further study.

### In vivo studies

All animal studies were conducted and monitored under an approved institutional and federal animal care guidelines and use committee (IACUC) protocol #201700322 at Emory University. For orthotopic experiments, 2 x 10^5^ murine PDAC cells were orthotopically implanted into the tail of the pancreas of 6–8-week-old female C57BL/6 mice (strain 000664, The Jackson Laboratory). After 10 days, tumor establishment was verified via bioluminescent imaging (BLI) using the In Vivo Imaging System (IVIS) (PerkinElmer #124262). Briefly, mice were injected with 180 𝜇L of 15 mg/mL D-luciferin (Biosynth #L-8220) dissolved in PBS and imaged using an open filter setting after 6 minutes. For model characterization and survival studies, animals were euthanized by CO_2_ asphyxiation followed by cervical dislocation upon reaching humane endpoint criteria. For adoptive cell transfer (ACT) therapy studies, mice were randomized such that animals in each treatment group had comparable tumor sizes. Subsequently, total body irradiation (TBI) was performed using 5 Gy of radiation from a SmART+ X-ray Irradiator (Precision X-ray) on day 10 post-surgery, followed by ACT therapy via tail vein injection 24 hours later (7.6 x 10^6^ mesoCAR+ T cells per mouse)^23^.

### T cell transduction

All experimental T cells were harvested from spleens of female B6.PL-*Thy1^a^/*CyJ (Strain 000406, The Jackson Laboratory) according to previously described protocols^24^. Lymphocytes derived from congenic mice were utilized for ACT therapy studies to distinguish donor CAR T cells (Thy1.1 (CD90)) from intrinsic host lymphocyte populations (Thy1.2 (CD90.2)). CD3^+^ T lymphocytes were negatively selected for using the EasySep™ Mouse T Cell Isolation Kit (StemCell Technologies #19851) and cultured in RPMI containing L-glutamine (Corning # MT10040CM) with 10 % heat-inactivated FBS (Atlas Biologicals #FS-0500-AD), 1 mM sodium pyruvate (Gibco #11360070), 1 mM HEPES (Gibco #15630080), 1 X MEM non-essential amino acids (Gibco #11140050), 0.55 mM β-mercaptoethanol (Gibco # 21985023), and penicillin-streptomycin (Corning #30-002-CI). αCD3/αCD28 magnetic beads (Dynabeads #11456D) were used according to manufacturer’s protocols and administered at a 1:1 bead to T cell ratio. One hundred I U/mL rhIL-2 (NIH repository) were added every 2 days, or as needed for 5 days, before initiating treatment. Best practices for IL-2 concentration, bead-activation, and culture conditions were based on recommendations in the literature^25^. MesoCAR-T cells were transduced with a chimeric anti-mesothelin single-chain variable fragment (scFV) fusion protein containing the T cell receptor CD3ζ signaling domain. CAR vector generation has been described for both human and mouse-specific mesothelin-binding scFvs^26,27^. The mesoCAR vector was generously provided by Dr. Carl June (University of Pennsylvania). CAR expression post-transduction was validated and quantified using a flow cytometry antibody specific to the human F(ab’)_2_ fragment (Jackson ImmunoResearch #109-606-006).

### Western blot

Proteins were extracted from cell lysates using mechanical dissociation via sonication, followed by suspension in RIPA lysis buffer containing 1% PMSF (Life Technologies #36978) and 1% phosphatase and protease inhibitor cocktail (Cell Signaling #5870). Protein concentration was determined using the Pierce™ BCA Protein Assay Kit (ThermoFisher #23227). Western blots were performed as previously described^28^. Primary antibody used to detect mesothelin was anti-mouse-mesothelin (Abcam# ab213174) and secondary antibody was anti-rabbit IgG, HRP-linked (Cell Signaling #7074). Anti-beta actin antibody was used as the loading control (Abcam #ab8226).

### Flow cytometry

Murine tumor tissue, spleen, and blood samples were collected for immunophenotypic analysis by flow cytometry. Organs were first mechanically digested by mincing tissue and then strained using a 70 𝜇m filter (Fisher scientific #08-771-2). Cell suspension was then chemically digested by incubation at 37°C for 30 min with collagenase (Roche #11088866001), dispase (Stemcell Technologies #07923) and liberase enzymes (Sigma Aldrich #5401119001). Single cell suspensions of each organ was incubated with antibodies (**Table 1**) in the dark at room temperature for 10 min, washed, and fixed in FACS Buffer containing 2 % FBS and 4 % para-formadehyde (Sigma-Aldrich #95-30525-89-4). Zombie Aqua Fixable viability dye (Biolegend # 423101) was used to detect live cells. Samples were measured on a Cytek Aurora flow cytometer (Cytek Biosciences). Between 10 x 10^4^ and 10 x 10^6^ events were collected per sample and analyzed using FlowJo software (Becton Dickinson v9).

**Table 1.**

| Flow cytometry antibodies | Company | Catalog | Concentration |
| --- | --- | --- | --- |
| Thy1.1 – PeCy7 | BD | 561404 | 1/500 |
| Thy1.2 – BV786 | BD | 564365 | 1/500 |
| Mesothelin – AF647 | Jackson ImmunoResearch | 109-606-006 | 1/500 |
| CD45 – BV570 | BioLegend | 103135 | 1/500 |
| CD3 – FITC | BioLegend | 100203 | 1/500 |
| CD4 – APC/Cy7 | BioLegend | 100413 | 1/500 |
| CD8a – V450 | BD | 560471 | 1/500 |
| CD26 – PE | BioLegend | 137803 | 1/500 |
| PD-1 – PerCP/Cy5.5 | BioLegend | 135207 | 1/500 |
| IHC antibodies |  |  |  |
| α-SMA | Abcam | ab7817 | 1/200 |
| CD4 | Abcam | ab183685 | 1/500 |
| CD8 | Abcam | ab237723 | 1/250 |
| Iba1 | Abcam | ab5076 | 1/200 |
| Thy1.1 | Biolegend | 202501 | 1/100 |

### Immunohistochemistry (IHC)

Formalin fixed paraffin embedded (FFPE) mouse tumors from *in vivo* mouse experiments were subject to IHC analysis as conducted by Emory’s Cancer Tissue Pathology Shared Resource. 4-5 µm slices were individually stained with Hematoxylin (Sigma #517-28-2), Picrosirius Red (Abcam #ab150681), and primary antibodies (**Table 1**) were developed using 3,3’-diaminobenzidin (DAB) substrate (Vector laboratories #SK-4100). Slides were scanned using an Olympus Nanozoomer whole slide scanner.

Images were analyzed using Qupath software (qupath.github.io). Total number of cells were quantified within a specified tissue area, and threshold detection was used to count the number of cells positive for a given antibody signal^29^. For picrosirius red, threshold detection was used to quantify positive area (µm^3^) of total tumor tissue area. Multiplex immunofluorescence staining included DAPI (Perkin Elmer #CS1-0127-2ML), CD4, and CD8 primary antibodies (**Table 1**), followed by Opal 690 and Opal 520-conjugated secondaries (Perkin Elmer), respectively. Images were acquired using a Roche BenchMark ULTRA IHC/ISH system autostainer.

### Gene expression analysis

Pancreatic and lung tumor gene expression and somatic mutation data was obtained from The Cancer Genome Atlas (TCGA) https://www.cancer.gov/tcga [PDAC^30^, LUAD^31^, LUSC^32^]. Pancreatic tumor cases were limited to the 150 samples previously established as ductal adenocarcinomas [PDAC^30^]. In total, 483 Lung Squamous Cell Carcinoma samples and 530 Lung Adenocarcinoma samples with both mutation and expression data were used. Mutation MAF files were used to identify samples with the TP53 R175H hotspot mutation. All other mutations were categorized as loss of function (LOF), and samples with other mutation data that included no mutation in the TP53 gene were considered wild-type. Expression values were log2 transformed, then the average was computed among established normal and activated stroma gene sets for each sample (https://github.com/rmoffitt/pdacR^33,34^). Comparisons between TP53 mutation status were preformed using Welch’s two sample t-test comparing average expression of loss of function group with hotspot group. Analysis was run in R version 4.2.1 with the stats package (R Core Team (2022). R: A language and environment for statistical computing. R Foundation for Statistical Computing, Vienna, Austria. Available online at https://www.R-project.org/.).

### Statistical analysis

Statistical analyses were performed on Prism 9 (GraphPad Software v9). Comparisons between two groups were done via student’s t-test, or if multiple groups were present, via one-way ANOVA, followed by pairwise t-test to compare groups for significance. *p<0.05, **p<0.005, ***p<0.0005.

## Results

### Syngeneic models differentially impact mouse survival and tumor composition

Orthotopic implantation of tumor cells into the pancreas of immune competent mice represents a practical approach for testing immunotherapy *in vivo*. We surmised that these models would recapitulate key stromal and immune features of the standard KPC-GEMM while permitting consistency in tumor burden. Survival studies were conducted to assess progression variability between distinct, KPC-GEMM-derived PDAC tumor lines post orthotopic implantation. KPC-Luc and MT-5 tumors were used for these studies **(Figure 1A)**. They are derived from two different KPC GEMMs, each driven by oncogenic Kras and TP53 alterations. TP53 alteration differed slightly between cells: the KPC-Luc cells harbor a null TP53 (TP53^-/-^) and MT-5 cells have a hot spot inactivating mutation (TP53^R172H/+^). Given these clinically relevant genetic profiles, we anticipated similar phenotypic properties of tumors would be evident upon implantation *in vivo*. Orthotopic KPC-Luc was a more aggressive model, with median survival of mice was 23 days MT-5-bearing mice survived a median time of 36 days **(Figure 1B)**. H&E staining revealed that MT-5 tumors closely mirrored the desmoplastic stroma and neoplastic glands present within KPC-GEMM tumors **(Figure 2A)**. In stark contrast, the KPC-Luc tumors displayed a spindle-like and sarcomatoid morphology, which were largely devoid of desmoplasia.

**Figure 1:**
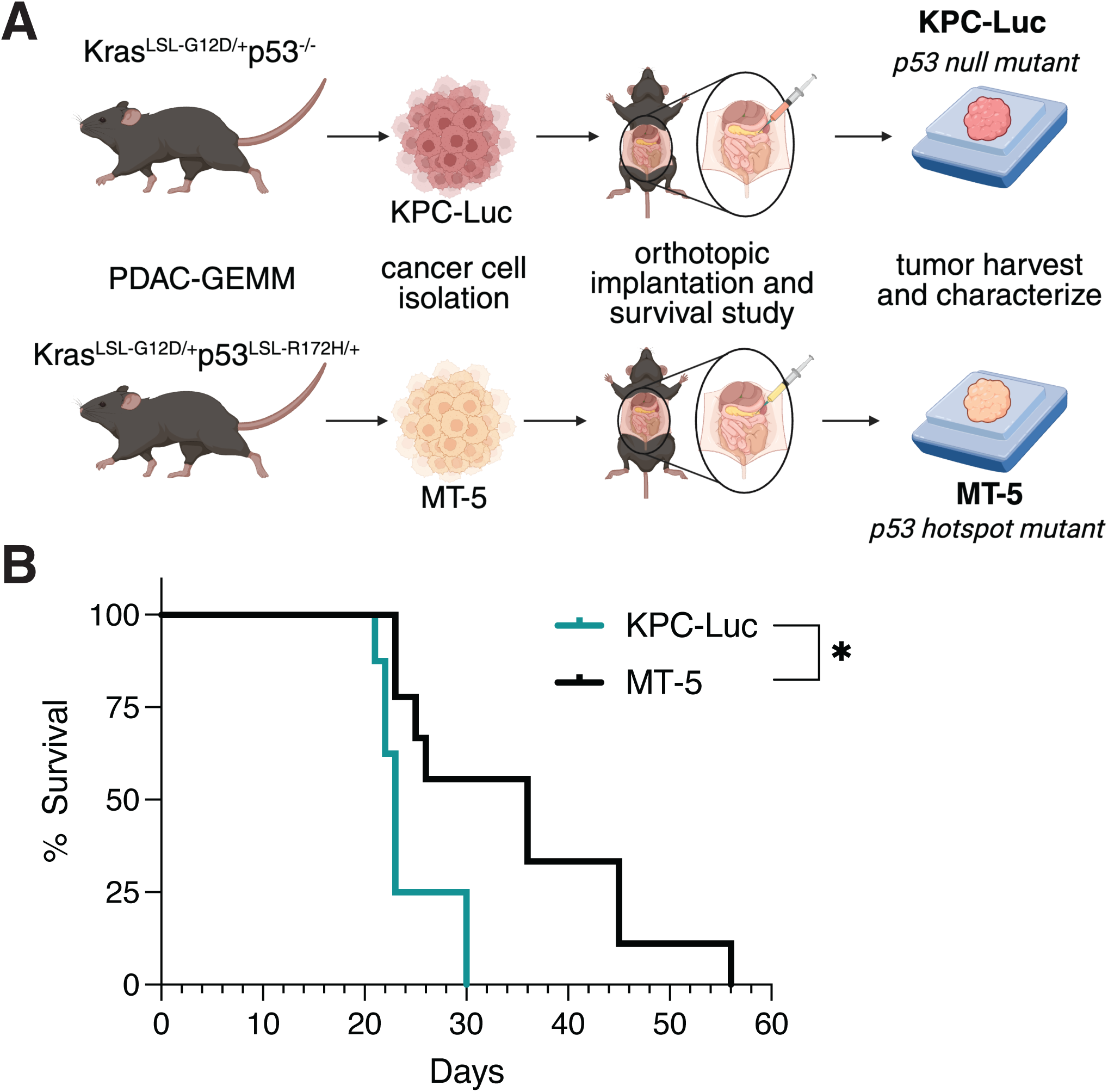
Orthotopic PDAC tumors recapitulate the TME of GEMM tumors. **(A)** Experimental schematic of cell isolation and orthotopic longitudinal survival study. Immortalized cell lines were isolated from KPC-GEMMs. **(B)** Survival kinetics of immunocompetent mice bearing KPC-Luc or MT-5 orthotopic PDAC tumors, n = 10 per group. One-way ANOVA = \**p*<0.05.

**Figure 2:**
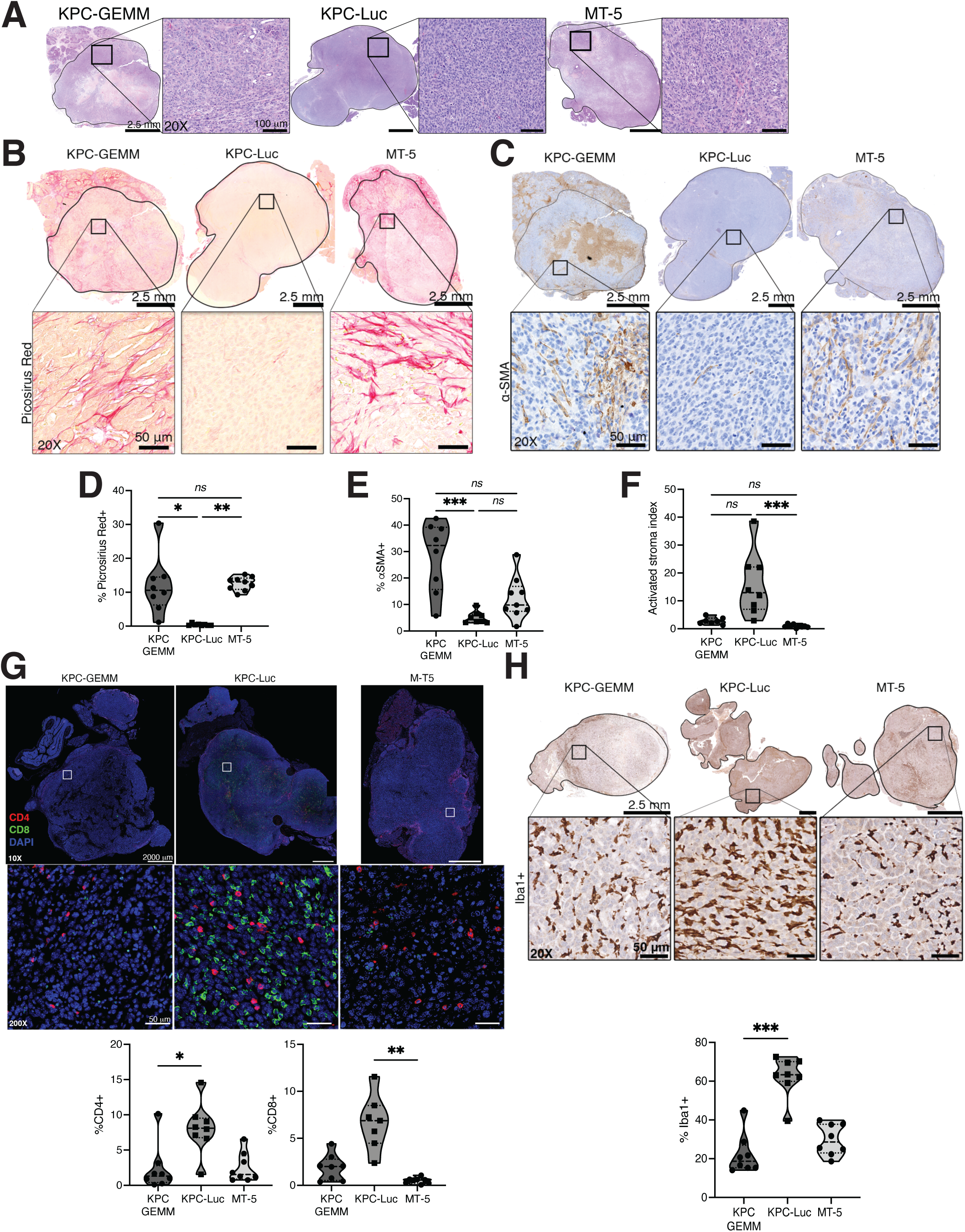
Orthotopic MT-5 PDAC tumors parallel the immune landscape of KPC-GEMM tumors. **(A)** Whole tissue (left) and 20X magnification (right) of H&E stained FFPE KPC-GEMM (left), KPC-Luc (middle) and MT-5 (right) tumor sections harvested at endpoint. Picosirus red **(B)** and alpha-smooth muscle actin (ɑ-SMA) **(C)** immunohistochemistry (IHC) staining of KPC-GEMM (left), KPC-Luc (middle) and MT-5 (right) FFPE tumor sections. Quantification of stromal fibrosis by picosirus red **(D)** and myofibroblastic positive stain by ɑSMA **(E)** as measured by percent (%) positive cells of total tumor area. The activated stroma index **(F)** was calculated by dividing values of ɑ-SMA quantification by picosirus red values. **(G)** Representative images of CD4+ (red) and CD8+ (green) T cell immunofluorescence on whole tissue sections of KPC-GEMM, KPC-Luc and MT-5 tumors. Quantification of CD4+ or CD8+ T cell infiltration (below) as measured by percent positive cells out of total cell count in tumor area. **(H)** Representative images of Iba1+ IHC staining on whole tissue sections of KPC-GEMM, KPC-Luc and MT-5 tumors. Quantification of Iba1+ positive IHC stain measured by percent positive cells out of total cell count in tumor. Kruskal-Wallis test = \**p*<0.05, \*\**p*<0.005, ***p<0.0005.

### Dense stroma correlates to reduced immune infiltration in orthotopic PDAC tumors

Stromal features were evaluated across the two orthotopic models and compared to established KPC-GEMM tumors from mice at 3-5 months of age via immunohistochemistry (IHC)^35^. We hypothesized distinct stromal features that influence disease progression would be clearly identified and limit access of lymphocytes into tumors. To assess this idea, the stromal composition was evaluated for markers of collagen (picrosirius red), myofibroblast cells (αSMA), and T lymphocyte infiltration (CD4 and CD8). Significantly more collagen was detected in MT-5 (*TP53 hotspot*) orthotopic tumors, as compared to tumors from mice bearing KPC-Luc (*TP53 null*) tumors **(Figure 2B and C**). αSMA was more prominent in MT-5 as compared to KPC-Luc tumors, although KPC-GEMM derived tumors had greater αSMA staining relative to the orthotopic models **(Figure 2D and E**). Stroma in MT-5 tumors more closely mirrored to that of KPC-GEMM-derived tumors **(Figure 2B-E**). An activated stroma index (ASI) (ratio of myofibroblast over collagen deposit^36^) exemplified this pattern, as KPC-Luc tumors had a high ASI compared to either MT-5 or KPC-GEMM tumors **(Figure 2F)**. An inverse relationship between these prominent stromal features and immune infiltrates was also evident, as CD4^+^ and CD8^+^ lymphocytes **(Figure 2G)** and Iba1^+^ macrophages **(Figure 2H)** were significantly elevated in KPC-Luc tumors and more sparse in MT-5 and KPC-GEMM tumors.

To facilitate longitudinal imaging for *in vivo* studies, the stroma-rich MT-5 cell line was transduced to stably express luciferase (MT-5-Luc). This was achieved by isolating single-cell colonies of MT-5-Luc to select for pure clonal populations that expressed the highest levels of luciferase but did not mediate immunologic rejection in animals upon orthotopic pancreatic implantation **(Supp. Figure 1A and B)**. Further, IHC analysis of αSMA^+^ myofibroblasts, collagen and CD4^+^/CD8^+^ lymphocyte infiltration indicated orthotopic MT-5-Luc tumors phenocopied that phenotype of parental MT-5 tumors **(Supp. Figure 1C and D)**.

We next explored the relationship between TP53 alteration and stromal gene signatures in human PDAC tumors using publicly available data from The Cancer Genome Atlas Program (TGCA)^30–32^. We hypothesized human PDAC tumors with hot spot TP53 mutations would show differential gene signatures of desmoplasia, as compared to those with wild type or loss of function (LOF) TP53. As the PDAC data set inherently contained very few patients with hot spot TP53 mutations, we expanded our evaluation by including lung adenocarcinoma (LUAD) and lung squamous cell carcinoma (LUSC), two cancer types driven by similar oncogenic signatures. Genomic datasets were organized according to TP53 mutational status (R172H, LOF, or wildtype) and assessed for signatures of normal or activated stroma, based on a previously published gene signature^33^ **(Figure 3A)**. Expression of stromal genes in human samples suggest (with limited statistical power, due to limited cases with hot spot mutations) a decrease in normal stroma (*p* = 0.05 for PDAC, 0.90 for LUAD, 0.02 for LUSC), with a corresponding more activated stroma (*p* = 0.96 for PDAC, 0.86 for LUAD, 0.23 for LUSC) **(Figure 3B-D)**. Taken together, the human data suggests an increase in cancer driven metaplasia and fibrosis in samples with the TP53 R172H hotspot mutation, compared to cases with other TP53 mutations. With limited numbers of cases with hotspot mutations available in TCGA, the trends lack power but anecdotally supports the observed trend in our mouse models **(Figure 3B)**.

**Figure 3:**
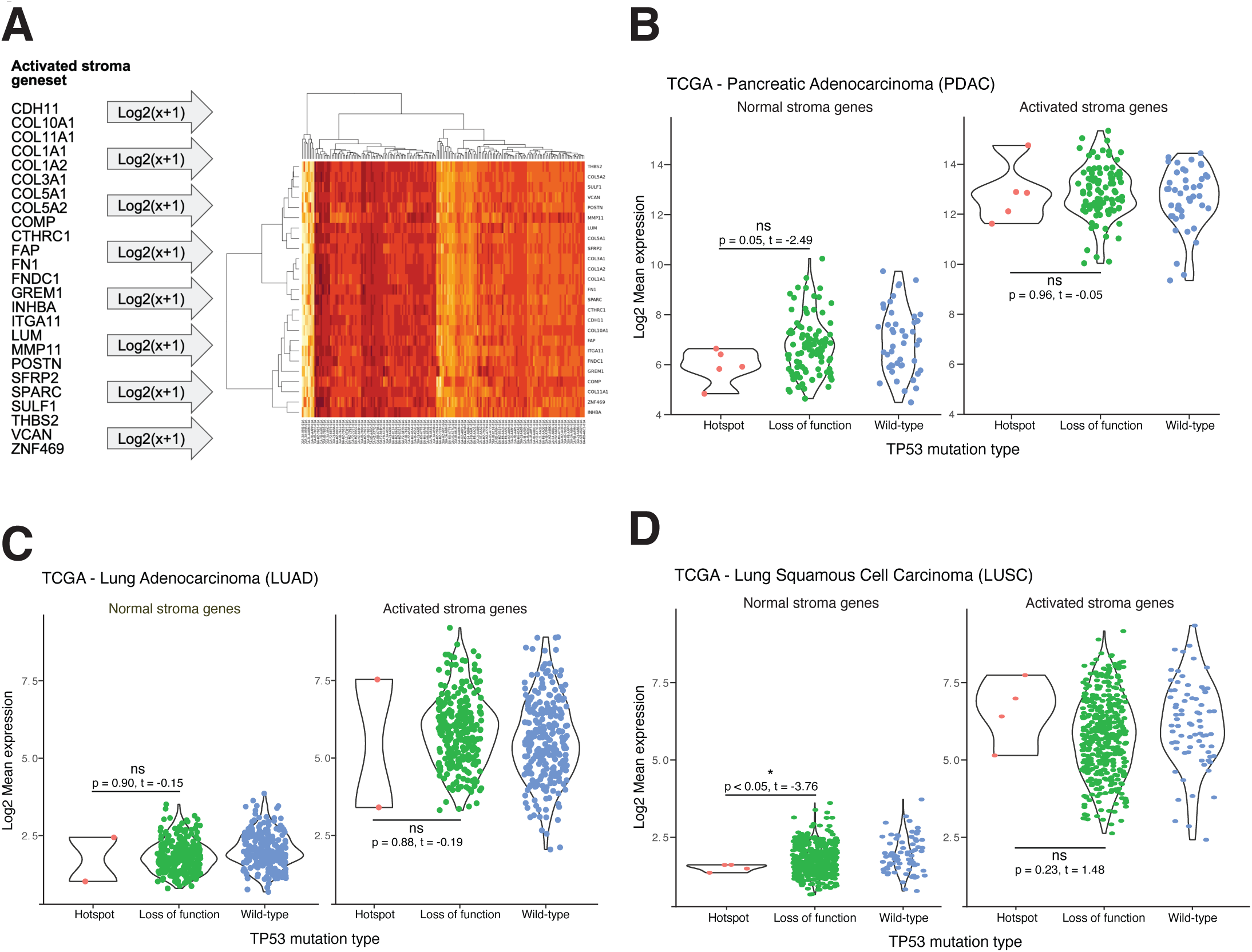
TP53 hotspot mutant tumors in humans contain activated stroma. **(A)** Heatmap stroma gene set and list of genes found in activated stroma from TGCA database. Comparison of activated stromal levels in human PDAC **(B),** LUAD **(C)**, and LUSC **(D)** tumors harboring TP53 hotspot (R175H) or Loss of function (LOF) mutations or not (wild-type) from TGCA human data.

### The desmoplasia of MT-5 tumors attenuates infiltration of MesoCAR T cells

We next tested our idea that tumors with more desmoplastic stroma blunt the ability of adoptively transferred T cells to infiltrate pancreatic malignancies. Past ACT approaches have evaluated the impact of stomal interference in PDAC mouse models by administering agents that interfere with integrity of the stroma or its associated components^37,38^. Yet studies on the efficacy of ACT across PDAC mouse models with varying levels of inherent stromal complexity are lacking. To address this idea in a fully immune competent system, bulk CD3^+^ murine T cells were engineered with a chimeric antigen receptor (CAR) directed against murine mesothelin (mesoCAR T cells) that signals CD3ζ and 41BB upon antigen recognition **(Supp. Figure 2A)**. The expression of this protein as an antigenic target was validated on both KPC-Luc and MT-5 murine PDAC cell lines. Immunoblot analysis was used to assess the presence of mesothelin as both a precursor and cleaved product (**Supp. Figure 2B**). Because CAR T cells are most adept at targeting cell-surface antigens, we further verified whether mesothelin was detectable on cell surface levels via flow cytometry by comparing protein levels to an unstained control (**Suppl. Figure 2C-D**). To address relationships between stroma and T cell trafficking, Meso CAR T cells were transferred to mice bearing bioluminescence imaged and verified, orthotopic KPC-Luc (stroma-poor, *TP53 null*) or MT-5-Luc (stroma-rich, *TP53 hotspot*) tumors. All Meso CAR T cells were administered 10 days following implantation, and one day following TBI as a lymphodepletion strategy **(Figure 4A**). On day 7, following ACT, mice were euthanized and analysis of Meso CAR infiltrating lymphocytes (TIL) by flow cytometry revealed significantly fewer present in stroma-rich MT-5-luc tumors, as compared to stroma-poor KPC-Luc tumors (**Figure 4B and Suppl. Figure 2E**). In contrast, Meso CAR T cells were readily detectable in blood of mice from both tumor models at equivalent levels (**Figure 4C**). Importantly, the abundance of donor Meso CAR T cells were validated in each tumor model via IHC staining for Thy1.1 protein levels (**Figure 4D - black arrows**) – a congenic marker found only on the donor MesoCAR T cells – and degree of desmoplasia at this time point was confirmed via IHC staining for picrosirius red (**Figure 4E**).

**Figure 4:**
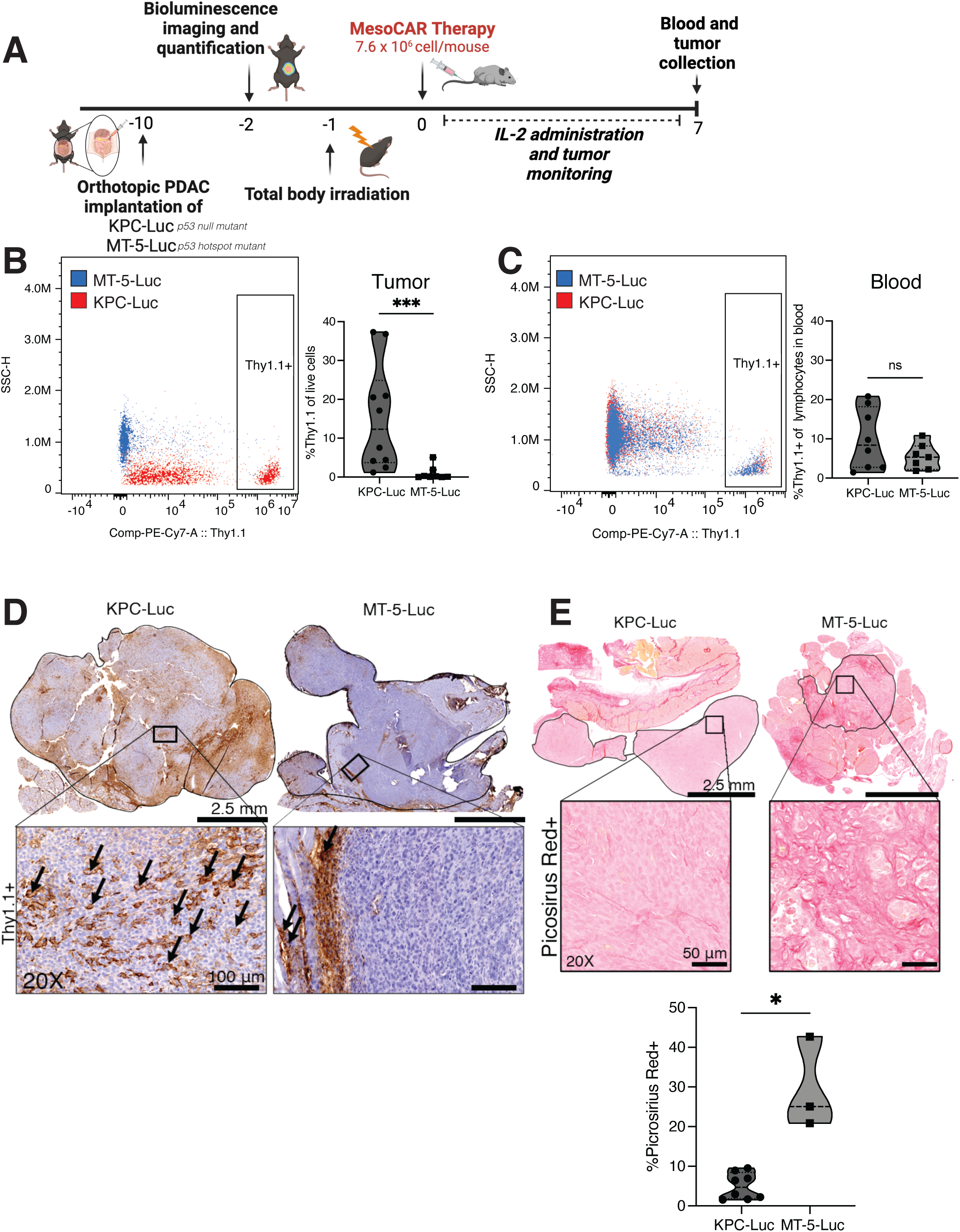
*TP53 hotspot mutant tumors mice block CAR T cell infiltration in mice.* **(A)** Study schematic. Mice were implanted with either KPC-Luc or MT-5-Luc PDAC tumors and after 10 days CD3+ mesoCAR T cell treatment was administered for 7 days. Tumors were harvested for either histological or flow cytometry analysis. Quantification of donor CAR T cells infiltrating tumor tissue **(B)** or circulating in blood **(C)** as measured by flow cytometry. Data represented as a percentage of Thy1.1 cells out of total live tumor cells (left) and as percentage of Thy1.1 cells out of total blood lymphocytes (right). **(D)** IHC representative images of Thy1.1+ stain in whole tissue and 20X magnification of KPC-Luc and MT-5-Luc FFPE tumor sections. Black arrows indicate positive Thy1.1 cells found in tumor parenchyma (left) or on tumor periphery (right). **(E)** Representative images of picrosirius red staining and quantification on KPC-Luc and MT-5-Luc FFPE tumor sections after CAR T cell administration, calculated by percent positive area of total tumor area. Mann-Whitney test = \**p*<0.05, ***p<0.0005.

## Discussion

The stroma of PDAC continues to be a formidable challenge for the immune system to overcome with the currently available cancer treatment modalities, highlighting the urgent need for new therapeutic avenues^16,17^. Previously published studies have attempted to enzymatically remodel the TME in combination with standard chemotherapy, but this approach did not improve objective tumor responses or overall survival. Current research suggests that higher quantities of tumor infiltrating lymphocytes (TILs) correlate with improved overall survival and progression free survival in patients with PDAC^19,20^. Therefore, ongoing efforts continue to focus on novel strategies to target components of the TME that increase lymphocyte infiltration. To do so, a representative model system that recapitulates the human malignancy and its abundant desmoplasia is vital. In many ways, GEMMs can parallel human PDAC by using similar oncogenic drivers and mutations in relevant tumor suppressors. GEMMs bearing these mutations develop spontaneous PDAC tumors that follow a similar path to malignant progression as human patients. Thus, GEMM tumors are often considered the “gold standard” for modeling tumor composition *in vivo*^39^. Unfortunately, GEMMs are highly variable, expensive, and time-consuming, creating significant drawbacks in their use for large animal studies. Due to the central role of the TME in PDAC progression and treatment outcomes, accurately modeling the microenvironment is crucial for reducing the disparity between preclinical data and human outcomes. By using immunocompetent, orthotopic tumor mouse models, our experiments have the dual-benefit of enabling donor-host immune interactions, as well as tissue site-specific tumor pathology that recapitulates the TME of a human PDAC patient. While human xenograft models lack reliable immune features and subcutaneous tumors lack site-specific physiology, our orthotopic model accommodates the best of both of these limitations.

Our data highlight both the utility and the limitations of distinct, immune-competent, orthotopic murine PDAC models for addressing hypotheses related to infiltration of therapeutic T cells. These results highlight unique phenotypic distinctions of the PDAC TME that occur despite seemingly similar genetic driver mutations in the tumor cells. Previous studies show that distinct TP53 mutations can drive differences in macrophage recruitment and lymphocytic populations of the TME^40^. Yet differences in desmoplasia in relation to mutational p53 burden in PDAC have not been documented and negatively correlated with infiltration of adoptively transferred CAR T cells to these tumors. By quantitatively comparing the composition of our orthotopic tumors to GEMM tumors, we show that the MT-5 model produces comparable TME features to those found in human PDAC patients. This finding is important, as it provides direct evidence for an inverse relationship between stroma and access of T cells that are targeted against relevant PDAC antigens. Mesothelin is over-expressed in human PDAC tumors and previous studies have validated its potential as a therapeutic target^20,41^. While targeting mesothelin offers a promising avenue for treating PDAC, evidence suggests it will not be sufficient on its own. In alignment with other reports^42^, these results support a role for desmoplastic stroma in disrupting access of tumor antigen-directed T cells into the TME and emphasize the importance of using realistic pre-clinical *in vivo* models to approximate clinical features of patient PDAC tumors.

Within the PDAC TME, collagens are the most abundant ECM proteins, and therefore largely contribute to desmoplasia^43^. A large body of literature describes the role of desmoplastic stroma in preventing drug and lymphocyte infiltration into PDAC tumors^17,18,44–46^. Studies modifying levels of TME collagen, by depleting subsets of CAFs, accelerated tumor growth and reduced animal survival ^37^. However, collagen within the TME, which frames and anchors tumor tissue structure, can also aid in tumor progression by reducing cancer spread and promoting CD8^+^ T cell infiltration^47^. By remodeling the PDAC stroma, pre-clinical studies indicate increased T cell infiltration and improved tumor control can be achieved through the use of TGFβ inhibitors in combination with ACT, or by targeting both stromal and tumor cells via a bispecific T-cell engager (BiTE)^38,48^. These data highlight a complex relationship between stroma composition and immune response in accordance with tumor development.

Our results support the notion that access of lymphocytes to PDAC tumors is limited by the cellular components of the TME and potentially the nature of mutations driving the malignancy^49^. By characterizing the differing stromal features of distinct orthotopic tumors, we can better adapt our pre-clinical testing of cellular or other immunotherapy approaches to maximize clinical relevance and applicability.

## Declarations

### Ethics approval and consent to participate

This study used anonymous human data and did not involve any direct interactions or interventions with human participants. As a result, formal ethics approval and informed consent were not applicable to this research.

### Consent for publication

All authors have reviewed and approved the final version of the manuscript submitted. We agree to the submission of this manuscript for peer review and potential publication. We confirm that this manuscript is an original work, and it has not been published elsewhere other than in poster presentations at scientific meetings, nor is it under consideration for publication in any other journal. A copy of the initially submitted version of this manuscript has been previously posted on medRxiv in an effort to disseminate this research.

### Availability of data and material

Gene expression and somatic mutation data was obtained from The Cancer Genome Atlas (TCGA) https://www.cancer.gov/tcga [PDAC^30^, LUAD^31^, LUSC^32^]. The dataset and code utilized in our research are available at https://github.com/rmoffitt/pdacR^33,34^.

### Authors’ contribution

Conceived and designed project: I.K., C.H., C.M.P., G.B.L.

Collected data: N.K.H.; I.K., M.P., M.W., J.H.

Contributed data or analysis: N.K.H., I.K., M.P., M.W., M.A.H., J.H., R.A.M.

Performed analysis: N.K.H., I.K., M.P., M.W., M.A.H., J.H., Z.M., R.A.M.

Wrote the paper: N.K.H., I.K., C.M.P., G.B.L.

## Acknowledgements

We want to acknowledge the cores at Winship Cancer Institute and Emory University that made this research possible including the Pediatric/Winship Flow Cytometry Core, Cancer Tissue Pathology Shared Resource and the Winship Cancer Animal Models Shared Resource*, under NIH/NCI award number P30CA138292.* The content is solely the responsibility of the authors and does not necessarily represent the official views of the National Institutes of Health. Figure schematics were created with BioRender.com.

## List of Abbreviations

ACT: adoptive cell therapy
ASI: activated stroma index
ɑSMA: alpha smooth muscle actin
CAF: cancer associated fibroblast
CAR: chimeric antigen receptor
DAB: 3,3’-diaminobenzidine
ECM: extracellular matrix
GEMM: genetically engineered mouse models
IHC: immunohistochemistry
KPC: Krax, p53, cre
LOF: loss of function
LUAD: lung adenocarcinoma
Luc: luciferase
LUSC: lung squamous cell carcinoma
mesoCAR: chimeric antigen receptor expressing T cells directed against mesothelin
PDAC: pancreatic adenocarcinoma
TBI: total body irradiation
TGCA: the cancer genome atlas program
TIL: tumor infiltrating lymphocytes
TME: tumor microenvironment

## Figure Legends

*Supp. Figure 1: MT-5-Luc cells do not undergo phenotypic changes.* **(A)** Luciferase signal intensity and absence of host rejection of MT-5-Luc tumors was confirmed via bioluminescence imaging (BLI) for 28 days. Tumor burden increased over time as measured by average radiance per region of interest (ROI). **(B)** Endpoint tumor weight (28 days) did not differ between MT-5- Luc and MT-5 tumor-bearing mice. **(D)** Quantification of desmoplastic stroma (via picosirus red histological staining) and T lymphocyte infiltration (via CD4 and CD8 IHC) showed no significant difference between MT-5 and MT-5-Luc. Mann-Whitney test.

*Supp. Figure 2: Generation of mesoCAR T cells and PDAC cells express high levels of mesothelin.* **(A)** Representative flow cytometry plots of CAR expression in MesoCAR-T cells and control (vehicle) cells post-transduction. **(B)** Representative western blot (left) of mesothelin precursor protein, cleaved mesothelin product and vinculin (loading control) levels from lysates of murine-derived PDAC cell lines (MT-5-Luc, MT-5 and KPC-Luc). Murine lung tissue lysate was used as a negative control. Quantification of protein levels (right) is represented as a ratio of mesothelin proteins (precursor or cleaved) to vinculin. **(C)** Cell-surface mesothelin levels on KPC-Luc, MT-5 and MT-5-Luc cell lines measured by flow cytometry. B16F10 melanoma cells were used as a negative control. Representative histogram of fluorescence intensity (left) and quantification relative to isotype levels (right). **(D)** Percent mesothelin positive cells relative to isotype-stained samples. **(E)** Representative flow cytometry gating strategy schematic used for data analysis shown in Figure 4B and C.

