## Supplementary figures and images for "Clinically relevant orthotopic pancreatic cancer models for adoptive T cell therapy"

### Supplemental figures

#### Supplemental Figure 1

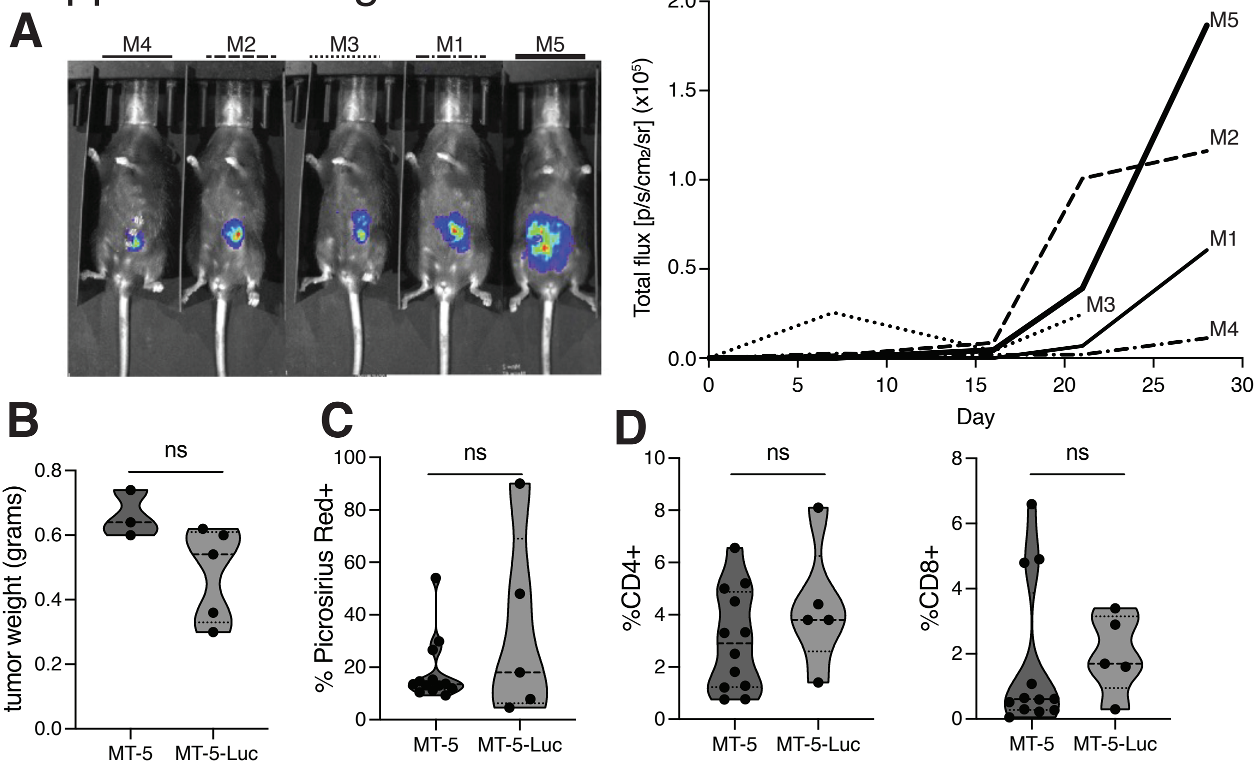

#### Supplemental Figure 2

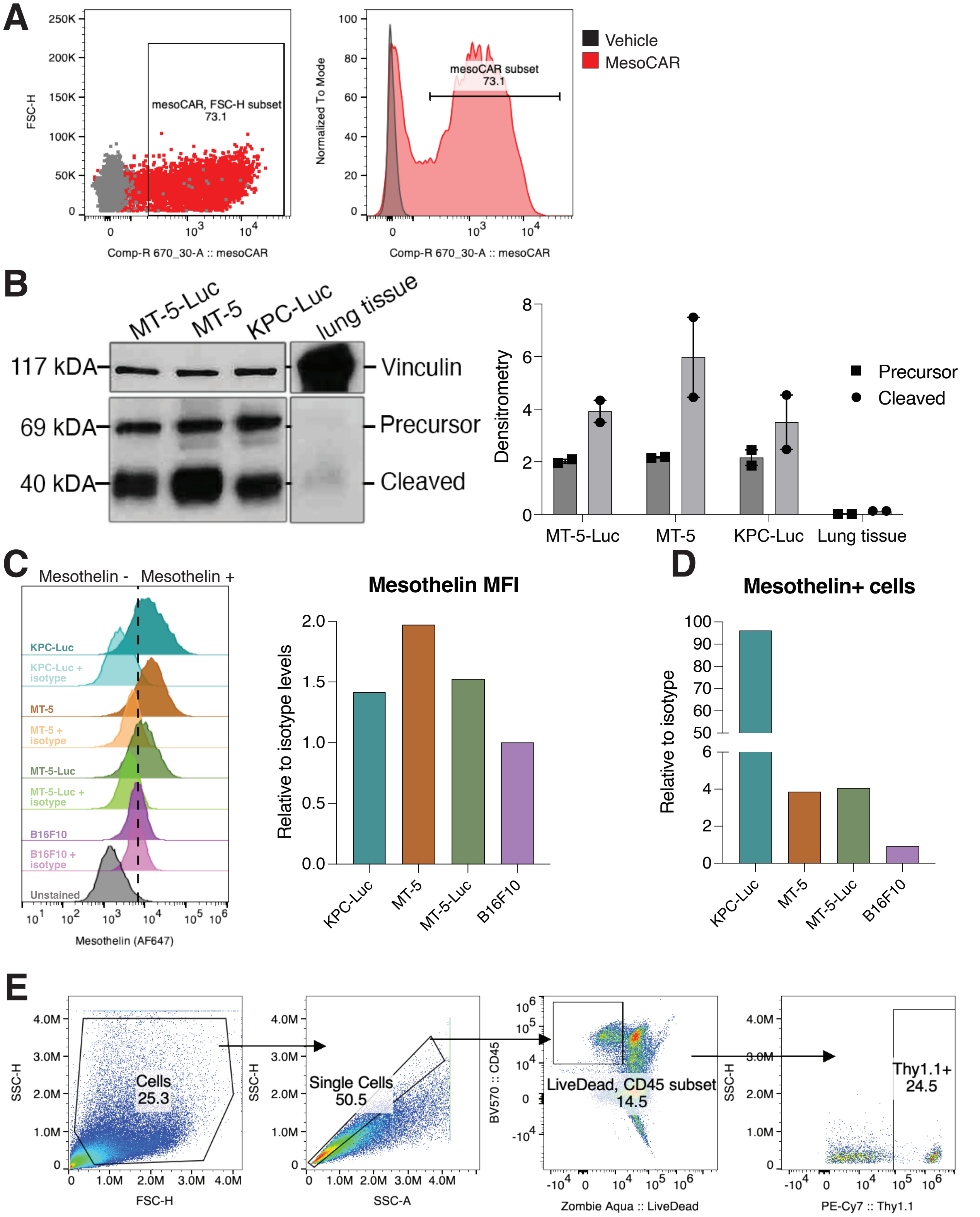
